# Selective increases in inter-individual variability in response to environmental enrichment

**DOI:** 10.1101/260455

**Authors:** Julia Körholz, Sara Zocher, Anna N. Grzyb, Benjamin Morisse, Alexandra Poetzsch, Fanny Ehret, Christopher Schmied, Gerd Kempermann

## Abstract

One manifestation of individualization is a progressively differential response of individuals to the non-shared components of the same environment. Individualization has practical implications in clinical setting, where subtle differences between patients are often decisive for the success of an intervention, yet there has been no suitable animal model to study its underlying biological mechanisms. Here we show that enriched environment (ENR) can serve as a model of brain individualization. We kept 40 isogenic mice for 3 months in ENR and compared the effects on a wide range of phenotypes on both mean and variance to an equally sized group of standard-housed control animals. While ENR influenced multiple parameters and restructured correlation patterns between them, it only increased differences among individuals in traits related to brain and behavior (adult hippocampal neurogenesis, motor cortex thickness, open field and object exploration, rotarod performance), in agreement with the hypothesis of a specific activity-dependent development of brain individuality.

## Introduction

Individualization is a process of developing unique traits and thus diverging from the inborn and genetically determined makeup. The behavioral and molecular bases of such divergence were traditionally investigated in human twin studies. However, the difficulty to conduct longitudinal studies in humans, as well as a limited range of phenotypes assessed in each twin cohort, leave many fundamental questions open. In particular, underlying mechanisms at the levels of cells, tissue, system or the entire brain and their interaction across the scales cannot be determined in human subjects due to the inability to collect all relevant phenotypes with sufficient depth and precision or to experimentally manipulate the processes in question. Thus, addressing these problems calls for a suitable animal model in which environment *and* genotype could be strictly controlled.

Individualization involves an increasingly differing response of initially highly similar individuals to exposure to seemingly the same environment. We propose that activity-dependent structural plasticity is a central mechanism contributing to the individualization of the brain. The iterative nature of the feedback loops between plasticity and behavior would result in increasingly different brains, behavioral trajectories and life courses. In this model, small initial differences are augmented through self-reinforcement. In support for this hypothesis, we previously showed that large groups of isogenic mice exposed to an enriched environment (ENR) developed stable and unique social and exploratory behavioral patterns that diverged between individuals over time (Freund et al., 2013; 2015). What differed between the mice of this cohort was their unique experience of that same environment and their resulting differential behavior. Because this ‘non-shared environment’ relates to the individual’s own experience and actions, the paradigm revealed a dimension that was previously largely hidden in group effects, but which is of greatest interest for studies addressing sources of variance in a system.

We and others had previously described the stimulatory effect of ENR on mean levels of adult hippocampal neurogenesis (Kempermann, Kuhn, & Gage, 1997; Nilsson, Perfilieva, Johansson, Orwar, & Eriksson, 1999; Tashiro, Makino, & Gage, 2007), the lifelong activity-dependent generation of granule cells in the mammalian dentate gyrus. Furthermore, we showed that longitudinal individual behavioral trajectories had correlated with the within-group differences in numbers of new neurons among the enriched mice, underpinning the suitability of adult neurogenesis as a biologically relevant readout of activity-dependent brain plasticity (Freund et al., 2013). While this previous experiment suggested an increased variance in numbers of new neurons integrated into the hippocampal circuit in ENR mice as compared to mice living in standard laboratory cages, the effect could not be unequivocally claimed due to the smaller size of the control group compared to the experimental group. Moreover, because behavioral assessment was based on monitoring animals in the ENR enclosure, the same constructs were not accessible for control mice. Finally, we could not conclude to which degree the effect of ENR on variance (and hence individuality) was specific to adult neurogenesis and exploratory behavior. The experiment to address these questions is presented here. Because ENR has been shown to influence a broad range of body and brain-related parameters, including metabolic states (Wei et al., 2015), volumes of certain brain areas (Diamond, Johnson, Protti, & Ott, 1985; Diamond, Krech, & Rosenzweig, 1964; Diamond et al., 1966), and different behavioral aspects (Clemenson et al., 2015; Garthe, Roeder, & Kempermann, 2015), it was of particular interest to test the ENR effect on the variance of these parameters. If increases in variance were general across all domains, this would suggest a common, non-specific causality. From a mechanistic perspective, the paradigm would thus be less feasible as a model to study the emergence of brain individuality. A more specific and selective induction of variance in response to enrichment would indicate that the observed individualization of the brain does not arise as a mere epiphenomenon of broader effects.

To investigate whether long-term environmental enrichment triggers the specific development of inter-individual differences between mice, we performed a cross-sectional study and analyzed differences in variance between groups of mice housed in one large enriched environment or in control cages (CTRL). Both ENR and CTRL groups consisted of 40 female C57BL/6 mice that were randomly assigned to their respective housing conditions, where they stayed for 105 days (Fig. 1A). In addition to the social complexity introduced by the number of animals in the enrichment cage, the complexity of ENR was increased by the large size and the compartmentalization of the enclosure (Fig. 1B). A total of 28 morphological, behavioral and metabolic variables were assessed (Suppl. Table S1).

**Fig. 1.**
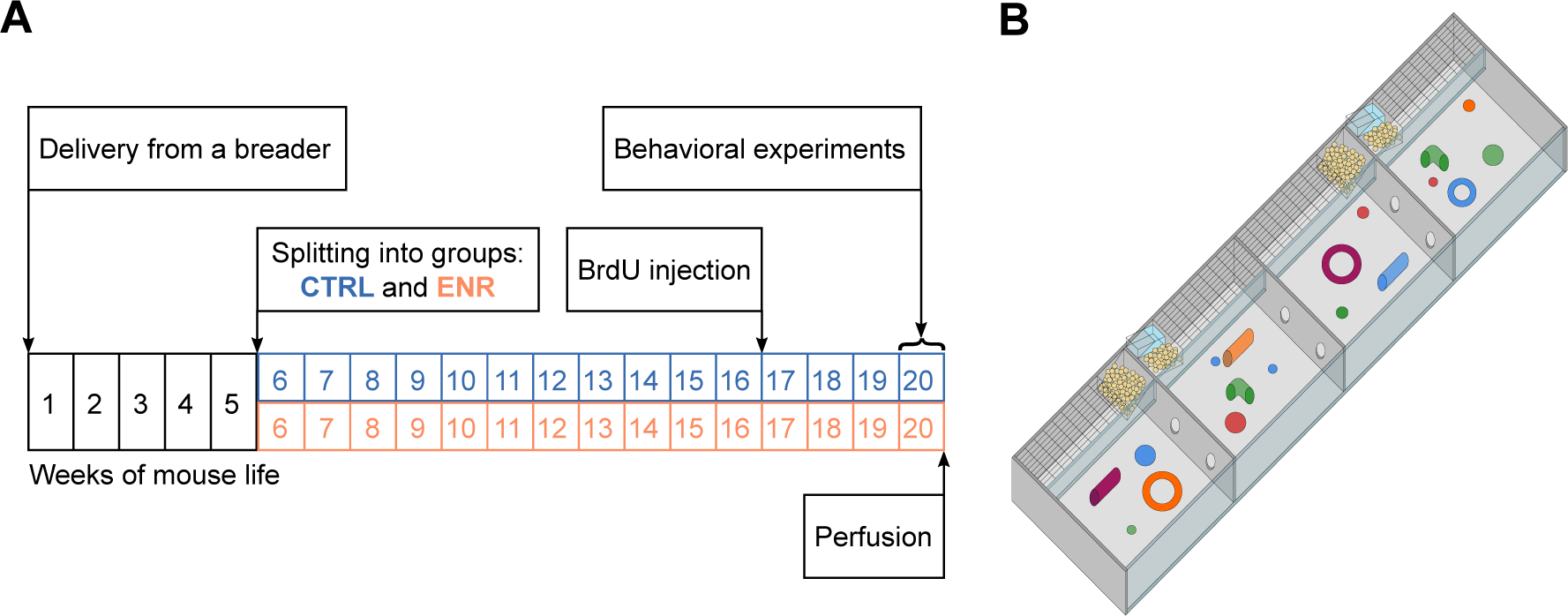
Experimental setup. (A) Experimental outline. At an age of 5 weeks, 80 female mice were split equally into two groups: one group lived in an enriched environment (ENR) for 15 weeks and one group lived in standard mouse cages in groups of five mice per cage (CTRL) for the same period of time. To analyze neurogenesis in the hippocampus, mice received intraperitoneal BrdU injections three weeks before perfusion. Behavioral phenotyping was performed in the last eight days before perfusion. (B) The enriched environment enclosure covered a total area of 2.2 m· and consisted of four sub-compartments, which were connected vial tunnels. Food, toys and nesting material were provided in every compartment.

## Results

### ENR reduces mean body size, but does not affect its variance

To determine the effects of ENR on gross body morphology, we monitored body weight of all animals over the course of the 105 days of the study (Fig. 2A). At the beginning of the experiment, no differences in weight existed between the two groups, confirming initial similarity between the randomized experimental mice. However, five weeks after the start of the experiment, ENR mice were significantly lighter than mice housed in control cages (CTRL). The difference in body weight remained constant throughout the experiment and indicated, together with the significantly shorter body length in ENR mice (Fig. 2B), that housing of mice in ENR reduces body size. In contrast, no differences in brain weights were detected between ENR and CTRL mice (Fig. 2C). The groups did not differ in the variances of body length, body weight and brain weight at any measured time point, suggesting that long-term ENR does not stimulate the development of inter-individual differences in gross body morphology.

**Fig. 2.**
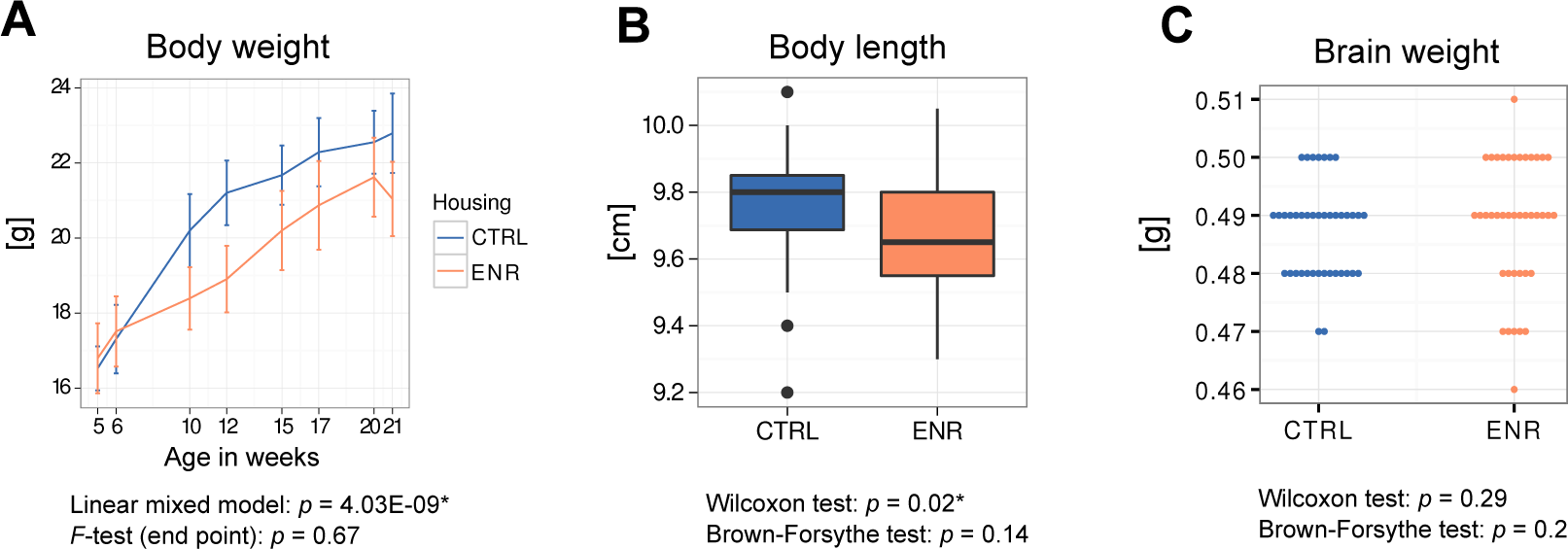
Environmental enrichment does not increase variance in gross body morphology. (A) Longitudinal measurement of body weight. Body length (B) and brain weight (C) were assessed at the end of the experiment.

### ENR increases inter-individual behavioral differences specifically in physical fitness and exploratory behavior

To analyze whether ENR increased inter-individual variability in behavior, all mice were subjected to the rotarod, the open field (OF) and the novel object recognition (NOR) tests (Fig. 3A-B). On the rotarod, ENR mice performed significantly better and showed a higher variance in their performance compared to controls (Fig. 3C), indicating that they developed individually different abilities for motor coordination, balance and mobility that were on average superior to CTRL mice. Despite the enhanced motor skills, ENR mice did not exhibit more locomotion during the OF and NOR tests (Fig. 3D-E), with the exception of the first OF trial, in which ENR animals traveled longer distances (Fig. 3D). In contrast, in the NOR test, enriched mice moved significantly less compared to CTRL animals (Fig. 3E). No significant differences were found in the variance of the distance animals traveled in any of the OF and NOR trials, suggesting that the increased inter-individual variability in the rotarod performance is not solely a consequence of variability in locomotion and broad activity levels.

In our previous work, we have introduced roaming entropy (RE) as a measure of territorial coverage and exploratory activity of mice in order to introduce a qualitative aspect into activity measurements (Freund et al., 2013; 2015). To investigate the effects of ENR on spatial exploration, we computed RE for all mice in the OF arena (Fig. 3F-G). On both days, ENR mice had significantly lower RE compared to CTRL animals. Moreover, on day 2, ENR mice showed a significantly greater variance in RE. Both ENR and CTRL animals habituated to the OF arena, which is indicated by a decrease in RE between the trials (Fig. 3H).

However, habituation was more pronounced in ENR mice. In the NOR test, ENR mice exhibited a significantly higher variance in the duration they explored the objects compared to CTRL mice (Fig. 3I), together indicating that ENR promoted the development of interindividual differences in interactions with the environment.

To examine the effect of ENR on recognition memory of the individual mice, we analyzed their abilities to discriminate the new object from the old one in the NOR test. Only a trend towards a higher preference for the new object was found in the ENR group compared to the CTRL group (Fig. 3J).

**Fig. 3.**
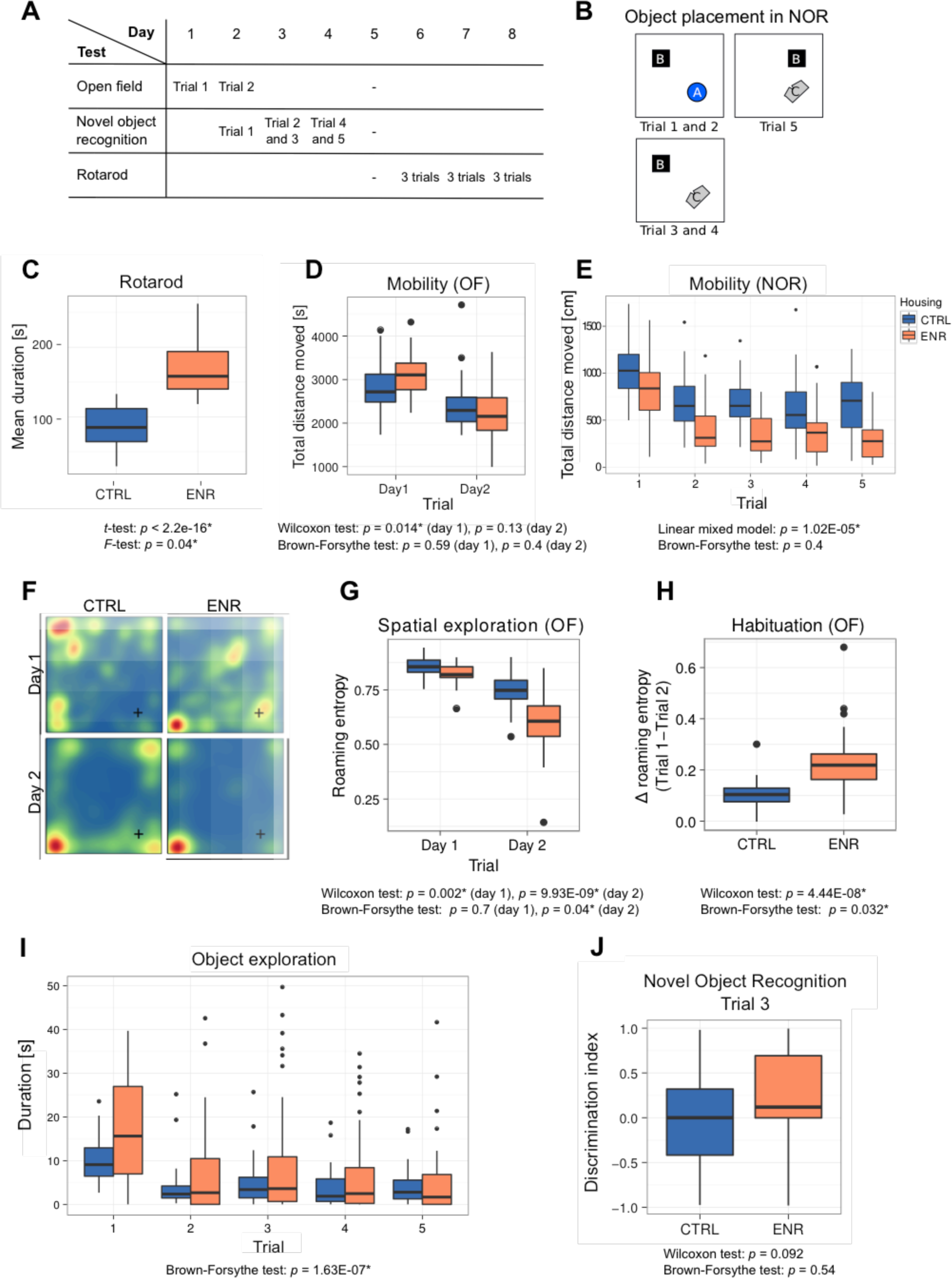
Mice living in an enriched environment exhibit inter-individual differences in motor abilities, spatial exploration and object exploration. (A) Timeline of behavioral testing. (B) Object placement in trials of the novel object recognition task (NOR). (C) Mean duration enriched mice (ENR, orange) and control animals (CTRL, blue) spent on the rotating rod in the rotarod task. (D) Total distance that mice moved in the arena on the two days of open field (OF) testing. (E) Total distance that mice moved during each trial in the NOR test. (F) Representative heatmaps for two mice depicting the probability of each mouse to be found at a specific location in the OF arena. Blue indicates lowest and red highest probabilities, respectively. The corner in which the light source was located is marked with a cross (+). (G) Roaming entropy in the OF arena describes spatial exploration. (H) Habituation to the OF expressed as a difference in roaming entropy between trials. (I) Object exploration in the NOR test. (J) Discrimination index indicating preference to the novel (+1) or the old object (−1).

### ENR increases inter-individual variability in the survival of new-born neurons

To assess whether the observed behavioral variability is reflected in differences in brain plasticity, we quantified the rates of adult neurogenesis in the dentate gyrus of the hippocampus. To estimate proliferation of precursor cells, we stained mouse brain sections for the proliferation marker Ki67 (Fig. 4A, E, F), whereas new-born cells that survived initial selection processes were identified by presence of BrdU, which was injected 3 weeks before the end of the experiment (Fig. 4B, G-I). No differences in mean or variance in the number of proliferating cells in the subgranular zone of the dentate gyrus were observed between ENR and CTRL mice (Fig. 4A). In contrast, we found a significant increase in mean and variance in the numbers of BrdU positive cells in animals housed in ENR (Fig. 4B), highlighting the specific effect of ENR on the survival of new-born cells. Co-localization of BrdU positive cells with the neuronal marker NeuN and the astrocytic marker 8100β showed that the variances in the survival of both cell types were higher in the ENR group compared to the CTRL animals (Fig. 4C-D). An increase in the total number of cells was, however, only found in the neuronal population. These results indicate that ENR increases inter-individual variability in the survival of new-born neurons and astrocytes but not in the proliferation of precursor cells in the dentate gyrus.

**Figure 4:**
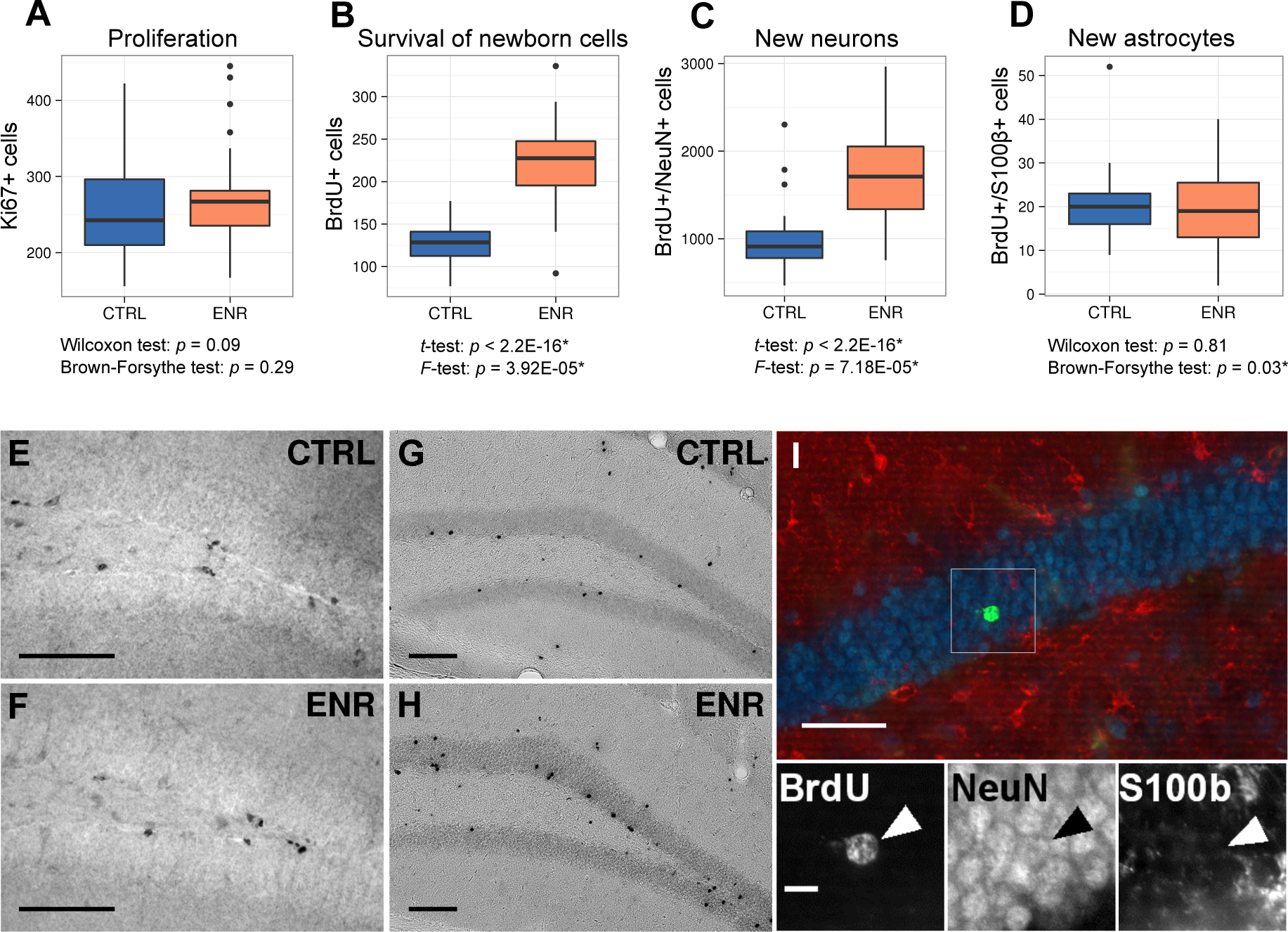
Environmental enrichment leads to the development of individual levels of adult hippocampal neurogenesis. (A) No difference in the number of proliferating (Ki67 positive) cells can be observed between mice housed under control conditions (CTRL) and mice housed in an enriched environment (ENR). (B) ENR mice have a significantly higher number and variance in the number of new-born cells as measured by quantification of BrdU positive cells three weeks after injection. The numbers of new-born neurons and astrocytes were determined by co-localization of BrdU with NeuN (C, I) and S 100β (D, I), respectively. Representative images of Ki67 (E, F) and BrdU immunostaining (G, H). (I) An optical section showing colocalization of BrdU positive cells (green) with NeuN (blue) and S 100β (red). The arrow highlights a new-born neuron. Scale bars are as follows: 100 *μ*m in E-H, 50 *μ*m in I, 10 *μ*m in I’.

### ENR does not elicit increases in variance of the hippocampus and cerebral cortex sizes

ENR has been long known to induce broad changes in brain structure, which lead to thickening of the cerebral cortex (Diamond et al., 1966), and increases in volume of the dentate gyrus (Kempermann et al., 1997). We also showed that the volume of the mossy fibers increases upon environmental stimulation concomitantly with adult neurogenesis (Römer et al., 2011). To further assess whether ENR increases inter-individual variability in brain plasticity beyond adult neurogenesis, we estimated the volumes of the hippocampus and its substructures: the dentate gyrus, infra- and suprapyramidal mossy fiber tracts (IMF and SMF) and the hilus. The volume of the entire hippocampus did not differ between ENR and CTRL mice (Fig. 5A). However, the dentate gyrus was significantly increased in the ENR mice (Fig. 5B). Furthermore, IMF (Fig. 5C) and the hilus (Fig. 5E), but not SMF (Fig. 5D), were significantly larger in ENR animals. None of these parameters showed different variance between housing conditions. This suggests that ENR differentially influenced various aspects of hippocampal plasticity, but the increased inter-individual variability triggered by ENR appears to be specific to adult neurogenesis.

Next, we measured the thickness of entorhinal, cingulate and motor cortex (Fig. 5M-O) since enrichment might specifically increase cortex thickness and structure in these areas (Diamond, 2001; Diamond et al., 1964). Although we did not detect differences in thickness of any of these cortices between CTRL and ENR mice (Fig. 5F-H), the motor cortex thickness showed a significantly higher variance in the ENR group (Fig. 5H).

**Fig. 5.**
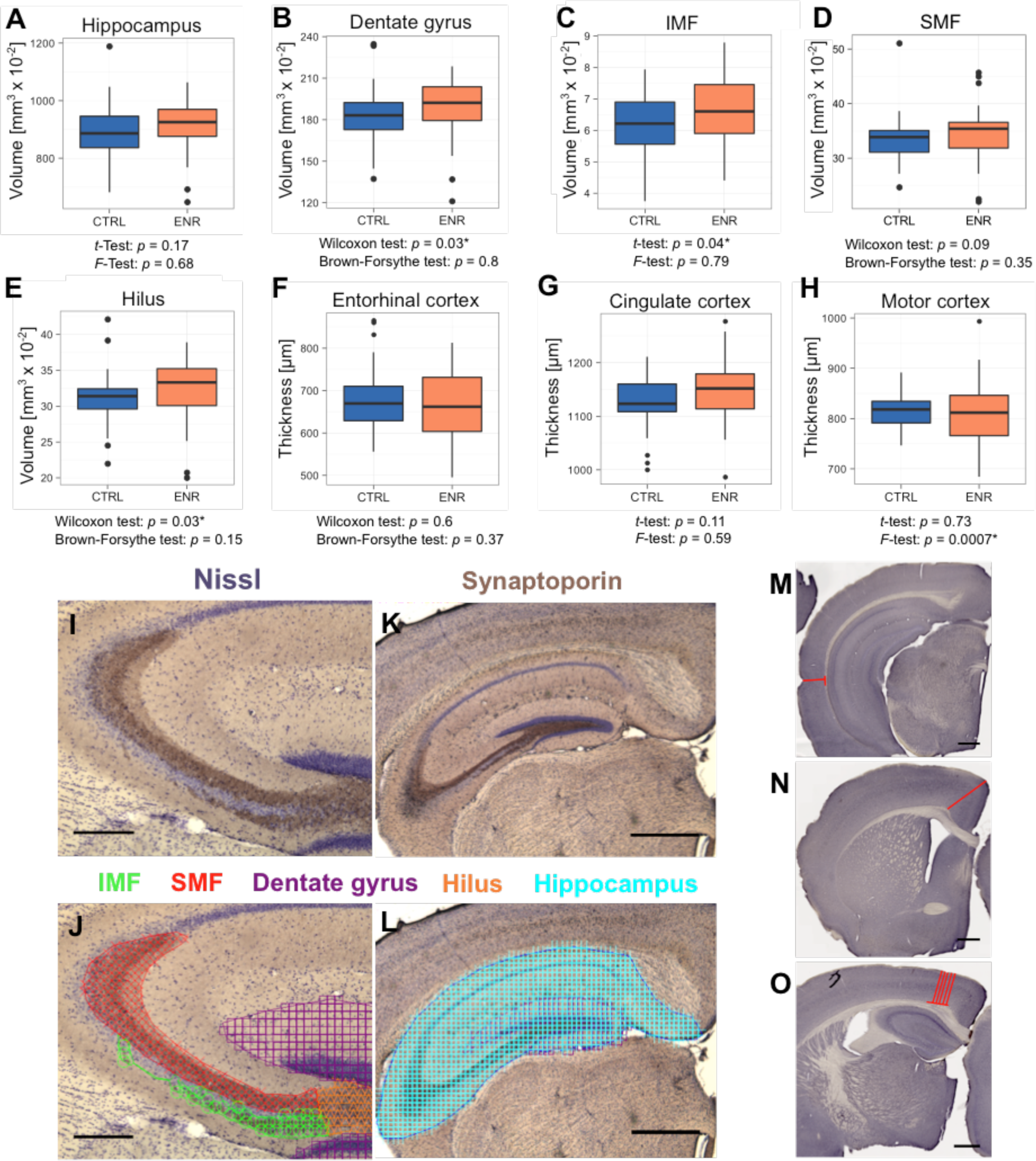
Environmental enrichment does not increase variability in gross brain morphology. (A-E) Volumetric analyses of the hippocampus (A), dentate gyrus (B), infrapyramidal mossy fiber tract (IMF; C), the suprapyramidal mossy fiber tract (SMF; D), and the hilus (E). (F-H) Thickness measurements of cortex areas, as indicated in the figure. (I-L) Representative images of sections immunostained with a synaptoporin antibody (brown) and counterstained with Nissl (purple). (J, L) Examples of contour tracing with overlaid Cavalieri probe estimator markers for the analyzed brain structures. (M-O) Thickness measurement on Ki67-DAB stained sections of entorhinal (M), cingulate (N) and motor cortex (O). Scale bars are 200 in I, J and 500 *μ*m in K-O.

### ENR reduces organ weights and cholesterol levels, but does not induce metabolic variability

Since ENR reportedly had beneficial effects on metabolism in outbred mice (Wei et al, 2015), we compared weights of liver and adrenal glands as organs playing a role in metabolic and hormonal regulation and analyzed basic blood biochemistry. In agreement with lower body weights, ENR animals had smaller adrenal glands and livers (Fig. 6A-B). The levels of plasma corticosterone, which is synthesized in the adrenal gland and is used as an indicator of animal stress, did not differ between the groups (Fig. 6C). Moreover, significantly lower plasma cholesterol levels (Fig. 6D) in ENR mice but no differences in glucose and triglyceride levels were seen between ENR and CTRL mice (Fig. 6E,F). No variance differences between the two groups were detected in any of the measured metabolic parameters.

**Fig. 6.**
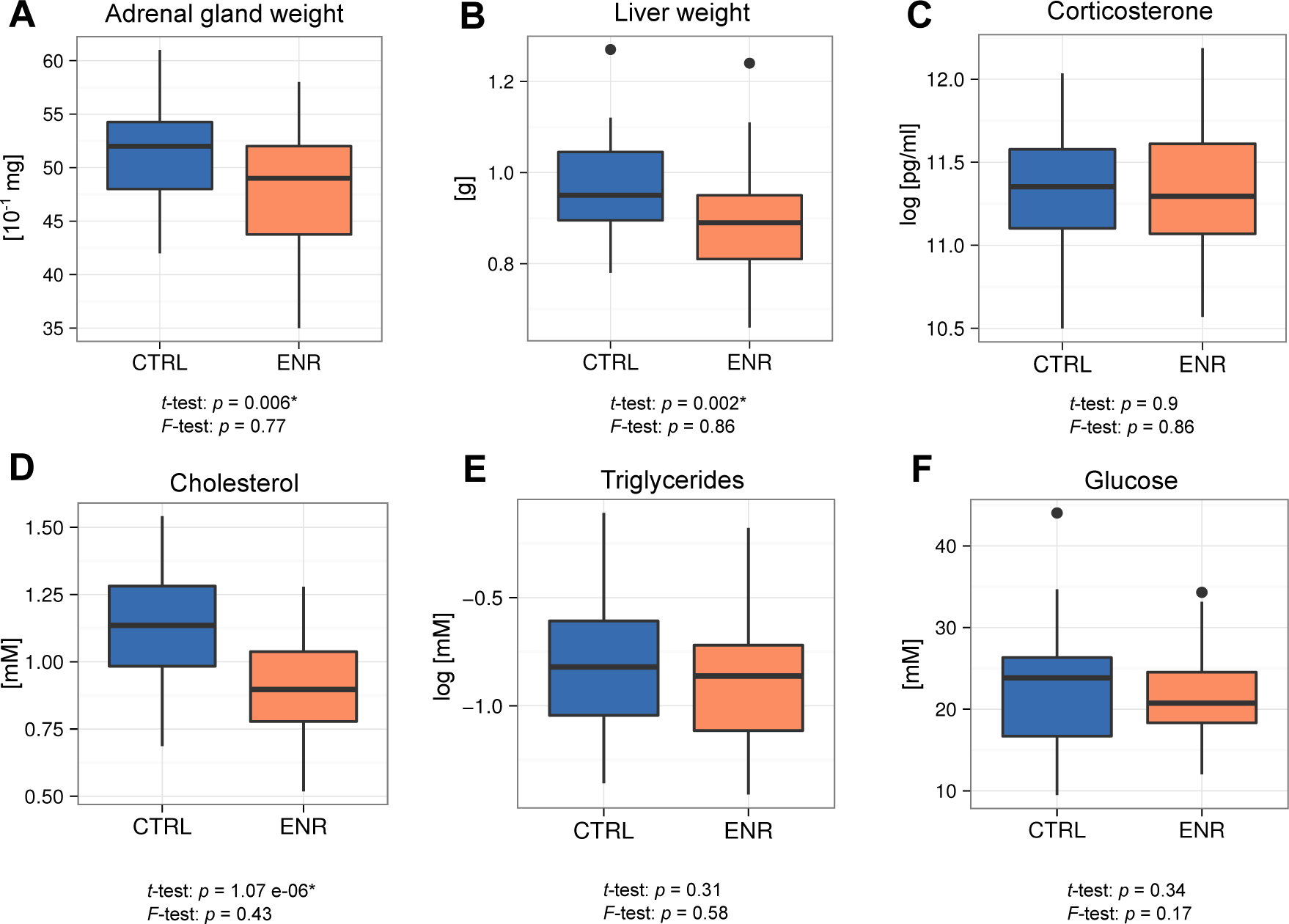
Environmental enrichment does not induce metabolic variability. (A, B) ENR decreases adrenal glands (A) and liver (B) weights. (C) Housing does not affect acute corticosterone levels. (D-F) Effects of ENR on plasma biomarkers.

### ENR restructures relationships between phenotypes

To analyze the impact of ENR on relationships between phenotypes we calculated correlations separately for CTRL and ENR mice (Fig. 7). Strong correlation between two traits would be pointing towards shared regulatory mechanisms controlling such parameters. We observed few significant correlations in either housing condition, which suggests that the majority of the traits were independent from each other. In both animal groups, we found significant positive correlations of roaming entropy and object exploration between trials of the OF and NOR tests, respectively (Fig. 7A). These correlations indicate that recorded behaviors were reliable and characteristic for each individual. ENR, however, weakened correlations between trials in NOR, as well as the negative association of object exploration to OF habituation, hinting towards more specific responses of animals to the environment (e.g. exposure to novel objects or their placement). Housing in ENR led to remodeling of associations between the brain structures and behavior (Fig. 7A). ENR uncoupled negative correlation of object exploration in NOR test as well as rotarod performance to the volume of the hippocampus. Similarly, habituation to the OF arena was negatively associated with the size of IMF and positively with the motor cortex thickness in CTRL, but not ENR mice. Hippocampal neurogenesis did not show significant correlation with any of the assayed phenotypes.

Metabolic phenotypes, namely plasma glucose, cholesterol and triglycerides were positively correlated in both housing conditions, but these relationships were weakened by ENR (Fig. 7B). As expected, plasma triglycerides correlated positively to the liver size in both groups. Epidemiological studies in humans suggest that brain and cognition are linked to the metabolism (Kapogiannis & Mattson, 2011; Panza et al., 2012). We observed few associations between the measured phenotypes in our mice (Fig. 7B). ENR changed the sign of the correlations between object exploration in NOR test and plasma cholesterol from positive to negative. It also promoted negative correlation between plasma glucose and rotarod performance, suggesting that fitness acquired by ENR mice has also a metabolic component.

**Fig. 7.**
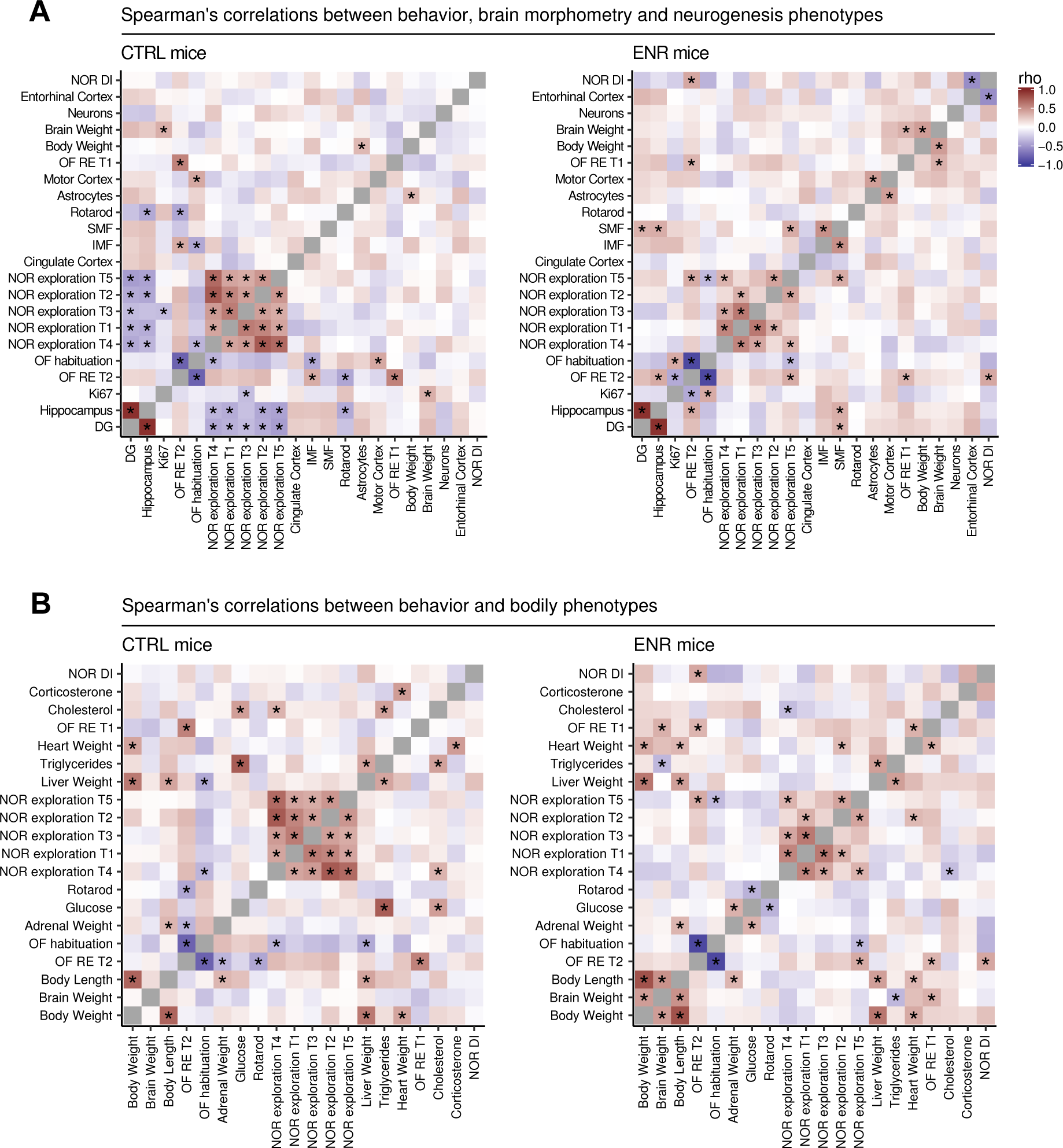
Environmental enrichment leads to partial restructuring of relationships between phenotypes. Heat maps show Spearman’s rank correlation coefficient between selected behavioral, morphometric and neurogenic (A) or metabolic traits (B) within the control (left panels) and the enriched (right panels) groups. Phenotypes were ordered based on hierarchical clustering in the ENR group. Significant correlations (*p* < 0.05) are marked with asterisks. NOR, novel object recognition test; OF, open field test; T1-5, trials 1-5 of the NOR and OF tests; RE, roaming entropy; DG, dentate gyrus; DI, discrimination index.

## Discussion

As medicine acknowledges inter-individual differences as a key determinant in diagnosis and treatment, understanding the biological mechanisms underlying individuality becomes increasingly important. Here we established that ENR is a suitable model to dissect processes leading to brain individualization. The purpose of our multivariate cross-sectional study of ENR effects in mice was to provide an insight into the magnitude of individualization of phenotypes spanning possibly broad aspects of physiology. We have shown that different traits are not uniformly affected by the stimuli, and, besides effects on the mean, for certain parameters, also effects on variance could be observed. These latter effects, in which enrichment increases differences between individuals in the group, cluster in variables related to behavior and adult neurogenesis.

Our behavioral data highlights a strong effect of ENR on animals’ active interaction with the environment: improved fitness and coordination, as assessed by the rotarod task; increased specific exploration of the environment, indicated by increased object exploration in NOR while the overall mobility decreased; and modification of general exploration and habituation, as demonstrated by the OF test. The effects of ENR housing on animal behavior in variations of OF and NOR tests have been reported in earlier studies. We have previously found, for example, that ENR mice habituate faster to an open field, which has been interpreted as improved spatial processing (Kempermann & Gage, 1999). Short 2-min trials in our NOR task precluded efficient familiarization with objects (Melani, Chelini, Cenni, & Berardi, 2017), which explains the lack of preference towards the new object. It has long been known that ENR elicits profound effects on brain plasticity and behavior (Mohammed et al., 2002; Sale, Berardi, & Maffei, 2014). The current study now highlights that ENR induced also substantial inter-individual variability in each of the assayed behavioral parameters. Particularly in NOR test, ENR mice showed much higher spread of object exploration times compared to the relatively homogenous CTRL group. Multi-trial design of the NOR and OF tests allowed us to confirm that the exploratory behaviors were stable and idiosyncratic, as indicated by the high intra-group correlations between the trials of each task. These observations corroborate our previous finding of ENR-induced individualization of spontaneous interactions with the environment (Freund et al., 2013). We have previously argued that the active participation with the outer world and habitat that manifests itself in the individual range of locomotion within that world (roaming entropy) is a major driving force of brain plasticity, presumably not limited to adult hippocampal neurogenesis (Freund et al., 2013. The current data is in agreement with this hypothesis. Improvement of rotarod performance in the ENR mice observed in this study implies that, even if the running wheels are not supplied, large cage area and toys to interact with provide considerable motor stimulation (Kempermann & Gage, 1999). Moreover, together with elevated variance of the motor cortex thickness, increased range of fitness in ENR mice point in the direction that ENR strongly affects plasticity of motor responses. It has been proposed that activation of motor cortex by ENR has widespread modulatory effects on other cortical areas (Sale et al.,2014; (Di Garbo, Mainardi, Chillemi, Maffei, & Caleo, 2011; Niell & Stryker, 2010).

Our previous report had suggested that long-term ENR induces variability in the survival of new-born neurons (Freund et al., 2013). The large size of both control and ENR groups in the present study now allowed us to corroborate that finding. To the degree that *p*-values can be taken as an indicator of effect strength (Nuzzo, 2014), the variance inducing effect of ENR on adult neurogenesis was the strongest among the measured phenotypes (*F*-test: *p* = 0.00004). Although new astrocytes did not increase in numbers, we also observed a small effect on variance. In contrast, there were no differences in variance in the number of proliferating cells between ENR and CTRL animals, even though there is a marginal effect of ENR on numbers of proliferating cells (*p* = 0.09), which suggests that over prolonged periods of time enrichment has subtle pro-proliferative effects (see also Ref. (Kempermann, Gast, & Gage, 2002)). Proliferating cells are a substrate on which selection mechanisms can act on, but their behavioral significance as such has not been shown so far. The fact that ENR does not trigger individuality in precursor cell proliferation indicates that individualization mechanisms act selectively only on those aspects of neurogenesis which are relevant for interactions with the environment. In contrast to the study by Freund et al., where 21% of variance in adult neurogenesis could be explained by differences in cumulative roaming entropy as an aggregated measure of the longitudinal behavioral trajectory, the current analysis detected no correlation between the number of newborn neurons and any of the cross-sectionally measured behaviors. The behavior in the familiar environment of the ENR cage could have been influenced by the group interactions (Shemesh et al., 2013), while the data presented here reflects the individual response of the animals to novel situations and might, therefore, constitute a different construct.

The mossy fiber projection, and especially its infra-pyramidal blade (IMF), is highly plastic (Crusio, Schwegler, & van Abeelen, 1989; Schwegler, Lipp, Van der Loos, & Buselmaier, 1981) and ENR can modulate its size (Römer et al., 2011). We here found an increased volume of the IMF but no effect on variance after 3 months of ENR, suggesting that even for aspects of hippocampal plasticity there is no simple parallelism in the effects of ENR. Similarly, we observed an increase in the volume of the dentate gyrus, but did not detect changes in variance or mean volume of the hippocampus. Adult-generated neurons contribute to the IMF (Römer et al., 2011), yet we did not observe a correlation between numbers of new neurons and IMF volume within either housing group, implicating that, under physiological conditions, other mechanisms than adult neurogenesis determine the bulk of the IMF. This finding is in agreement with the results from the screen in the genetic reference population (Römer et al., 2011).

Cortex thickness changes upon enrichment were reported in the older literature and have been the corner stone of the growing impact of the ENR paradigm in the 1970s (reviewed in Refs (Diamond, 2001; Rosenzweig & Bennett, 1996)). While increases in cortical thickness do not strictly mirror volume changes (Hammelrath et al., 2016; Winkler et al., 2010), they are an indication of massive cellular rearrangements in the cerebral cortex (Diamond et al., 1964; 1966). In our study, the only effect we found was an increase in the variance of the motor cortex thickness induced by ENR. The key difference to the old reports has been that we worked with mice, whereas essentially all classical studies had been done in rats. Indeed, the dynamics of three-dimensional brain development during first months of life is different between these two species (Hammelrath et al., 2016). Furthermore, the majority of old experiments compared enriched animals to impoverished littermates, which were kept in social isolation. Such impoverishment negatively affects brain size (Fabricius, Helboe, Steiniger-Brach, Fink-Jensen, & Pakkenberg, 2010), thus amplifying relative effects of enrichment (Bennett, Diamond, Krech, & Rosenzweig, 1964). We believe that the impact of ENR on cortical plasticity deserves still more specific analyses with much greater resolution.

It had been shown that ENR influences metabolism (Wei et al., 2015): keeping outbred mice in ENR resulted in decreased body weight, mostly through reducing fat content; lowered blood cholesterol, triglycerides, and glucose; and improved insulin and leptin signaling. It has to be noted that cages in the experiments performed by Wei and colleagues were equipped with running wheels to stimulate physical exercise of the animals. In the present study, we also observed decreases in body and liver weights, as well as lowering of plasma cholesterol, which indicates that ENR alone has a moderate beneficial effect on metabolism even in the absence of intense physical exercise. ENR animals also had smaller adrenal glands, and even though corticosterone levels were similar, this points towards reduced stress in ENR mice compared to CTRL. Finally, we did not detect differences in variances in any of the bodily parameters, further substantiating the conclusion that individualization of behavior and brain plasticity by ENR is not an epiphenomenon of more global physiological divergence.

Although the issue of variance has been brought up in very early studies (Walsh & Cummins, 1979), the ENR literature has not been concerned much with variance effects and interindividual differences. The focus has always been on mean group effects. The question of ENR effects on variance came up, however in the context of a movement in animal husbandry to provide larger space and enriching cage accessories in order to improve animal wellbeing and provide more species-appropriate conditions. Variability induced by ENR, the concern went, would work against the desired standardization and stability of animal experiments in the life sciences. A widely cited study by Wolfer et al., however, confirmed that ENR “increases neither individual variability in behavioral tests nor the risk of obtaining conflicting data in replicate studies” (Wolfer et al., 2004). The results presented here (Fig. 3) stand in clear contrast to the first part of this statement and potentially also the second. As we did not test a full spectrum of behavioral tasks, we must not generalize our conclusion beyond rotarod, open field and novel object recognition tests (this study), or free roaming in the cage (Freund et al., 2013). We would hypothesize that behavioral traits related to exploration and adjusting to novel situations, including hippocampal learning, are more strongly affected than others. The conclusion from Wolfer et al. requires a careful qualification. We nevertheless fully agree with the overall conclusion that “housing conditions of laboratory mice can be markedly improved without affecting the standardization of results”, especially if group sizes are sufficiently high. For most variables, even 3 months of ENR did not increase variability or alter correlations with other phenotypes. Furthermore, systemic variation might actually improve reproducibility (Richter et al., 2011) (see also ref. (Richter, Garner, Auer, Kunert, & Würbel, 2010) with comments and re-analysis in ref. (Jonker, Guenther, Engqvist, & Schmoll, 2013) and ref. (Wolfinger, 2013)). And finally, the attempt to ignore the within-group variation as an expression of a differential response to the same nominal stimulus might actually contribute to the “reproducibility crisis” to a much larger extent than previously appreciated.

Despite the ample literature on ENR, few studies addressing larger numbers of dependent variables have been conducted, and to our knowledge, we are first to investigate the interactions between an extended panel of variables in a correlation matrix. Similarly, there has been little insight into the isometry or allometry of the induced changes. Because our experiment employed large groups of animals, we could survey the inter-individual correlation patterns between the variables separately within each environmental condition, thus avoiding spurious relationships that could arise from mean differences between groups. Correlation matrices revealed the extensive relative independence of outcome measures, which suggests that the choice of traits for the analysis was broad enough to reflect distinct underlying causalities. Furthermore, ENR restructured correlation patterns by strengthening or weakening some associations (for details see Fig. 7), further demonstrating uneven regulatory influence of ENR on various aspects of physiology and plasticity. Thus, even in the absence of global mean effects on these parameters, ENR seemed to induce broad adaptations in brain plasticity and metabolism.

In conclusion, ENR does not generally increase variability. Increased variance in physical fitness (rotarod), exploratory behavior, adult neurogenesis and motor cortex stand out in the net size of the variance-enhancing effect. Their correlation pattern with other ENR effects is complex, though, with ENR remodeling many of the associations. ENR arises from this study as a more holistic paradigm than often assumed and proves to be a tool of choice to investigate the bases of individualization. In the laboratory setting, animals are relieved from pressures present in nature and therefore they are free to choose the degree of interaction with the environment. In our previous longitudinal study, we made the case that in a situation, in which both genes and (nominal) environment are kept constant, individuality emerges as a consequence of the so-called “non-shared environment”, i.e. the individual response to that environment and activity (Loseva, Yuan, & Karnup, 2011). Our current cross-sectional data suggests that multivariate studies with a large number of individuals and, ideally, a longitudinal design are needed to elucidate the exact contribution of the non-shared environment to the overall outcome of increased individualization.

## Materials and Methods

### Animal husbandry

80 female C57BL/6J mice were purchased from Janvier at an age of 4 weeks and housed in standard polycarbonate cages (Type III, Tecniplast) in groups of 5 until the start of the experiment (Fig. 1A). At an age of 5 weeks, 40 mice were randomly selected and transferred into the enriched environment, where they stayed for three months. The enriched environment consisted of four quadratic polycarbonate cages (0.74 × 0.74 m) that were assembled in a row and connected by two plastic tubes each. In total, the enriched environment covered an area of 2.19 m^2^ (Fig. 1B). Food and water were provided in every compartment of the cage. Animals in the enriched environment did not receive special food. To provide sensory stimulation, each compartment of the cage was equipped with plastic toys, tunnels and hideouts, which were cleaned and rearranged once per week. The bedding material was replaced on a weekly basis. Once a month, the entire enclosure was cleaned. Control animals were housed for the same period of time in standard polycarbonate cages in groups of 5. All mice were maintained on a 12-/12-hr light/dark cycle with food and water provided *ad libitum.* Three weeks before sacrifice, mice were injected intraperitoneally with bromodeoxyuridine (BrdU; 50 mg/kg body weight; dissolved in 0.9% NaCl). Injections were performed once per day for three consecutive days. All experiments were conducted in accordance with the applicable European and national regulations (Tierschutzgesetz) and were approved by the responsible authority (Landesdirektion Sachsen).

### Behavioral tests

Before starting the behavioral experiments, every mouse was visibly marked at the tail. To simplify handling, enriched animals were placed in the morning of every test session into standard cages in groups of five, which remained consistent throughout testing, and returned into the enriched environment cage in the evening. Mice were tested in the same order in all behavioral tasks. The sequence of the behavioral experiments is shown in Fig. 3A.

### Rotarod

Mice were assessed for locomotor abilities using an Economex Rotarod from Columbus Instruments. The rotating cylinder started with a speed of 4 rpm and accelerated by 0.1 rpm. At a final speed of 34 rpm and a maximum time of five minutes, the test was stopped manually. The trial was completed when an animal fell off or reached the maximum duration. The mice were trained on three consecutive days with three trials per day. The rotarod was cleaned after every session.

### Open field test

The open field (OF) enclosure consisted of a 120 × 120 cm squared apparatus subdivided into four identical arenas of 60 × 60 cm, allowing for the simultaneous testing of four mice in the apparatus. The 40 cm high white plywood walls were marked with a green tape on the intersections to provide additional spatial clues. The only light source in the room, a 100 watts light bulb, was installed 1.5 m above the intersection of the middle walls, next to the camera (Logitech). Mice were placed in the middle of the empty arena and were allowed to freely explore the arena for 5 min in each trial. The total of two trials were performed on two consecutive days. Roaming entropy (RE), a measure of territorial coverage, was calculated according to (Freund et al., 2013). Each arena was divided into 10 × 10 subfields. Probability *p*_*i*_ of a mouse being in a subfield *i* was estimated as a proportion of trial time spent in that subfield. Shannon entropy of the roaming distribution was then calculated as:

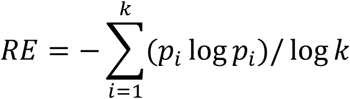

where *k* is the number of subfields in the arena (*k* = 100). Dividing the entropy by the factor log(k) scales the RE to the range from zero to one. RE is minimal for the mice that stay in one place and maximal for the mice that spent equal amount of time in each subfield of the arena. Data from 8 CTRL animals was lost in the second OF trial.

### Novel object recognition test

The two OF trials are to be understood as habituation for the NOR task (Fig. 3A). For this task, the same arenas were equipped with two different objects: object A was a 3.5 cm high blue cylinder with a diameter of 1.5 cm, object B was a black box of 8.5 × 9.5 × 2 cm, whereas object C was 4.5 cm long and transparent with a more complex geometric shape. For object placement in subsequent trials, see Fig. 3B. On day 1, following the OF trial, mice were presented objects A and B. On day 2, animals were first exposed to the same objects and in the following trial object A was replaced with object C. The same combination of objects was presented on day 3, followed by a trial in which object B was moved into the adjacent quarter of the arena. Each trial lasted 2 min. Discrimination index was calculated for trial 3 based on the exploration time for the new and old object as follows: DI = (new object - old object)/(new object + old object), and ranged from −1 (preference for the old object) to 1 (preference for the new object), while 0 indicated no preference (Miyauchi, Neugebauer, Oyamada, & Meltzer, 2016). 11 ENR and 3 CTRL mice, which did not explore object A in any of the first two trials, or did not explore any object in the trial 3 were excluded from the calculation of DI.

### Tissue preparation

Mice were deeply anesthetized with a mixture of ketamine and xylazine and transcardially perfused with 0.9% NaCl. Directly after the perfusion, liver, heart and adrenal glands were harvested and weighed. Brains were removed from the skull and postfixed in 4% paraformaldehyde overnight at 4 °C and equilibrated with 30% sucrose in phosphate buffered saline (PBS). For immunohistochemistry, brains were cut into 40 *µ*m coronal sections using a sliding microtome (Leica, SM2000R) and stored at 4 °C in cryoprotectant solution (25 % ethyleneglycol, 25 % glycerol in 0.1 M phosphate buffer, pH 7.4).

### Immunohistochemistry

For detection of BrdU, Ki67 and synaptoporin positive cells, immunohistochemistry was performed using the peroxidase method as previously described (Steiner at al, 2008). Briefly, free-floating sections were incubated in 0.6% H_2_O_2_ for 30 min to inhibit endogenous peroxidase activity. After washing, non-specific antibody binding sites were blocked using 10% donkey serum and 0.2% Triton-X100 in Tris buffered saline (TBS) for 1 h at room temperature. For BrdU detection, prior to blocking, sections were incubated in prewarmed 2.5 M HCl for 30 min at 37°C, followed by extensive washes. Primary antibodies were applied overnight at 4 °C as follows: monoclonal rat anti-BrdU (1:500, Serotec), rabbit anti- Ki67 (Novocastra, 1:500), rabbit anti-Synaptoporin (Synaptic Systems, 1:500). Sections were incubated with biotinylated secondary antibodies for 2 h at room temperature (1:500, Dianova). Primary and secondary antibodies were diluted in TBS supplemented with 3% donkey serum and 0.2% Triton-X100. Detection was performed using the Vectastain ABC-Elite reagent (9 μg/ml of each component, Vector Laboratories, LINARIS) with diaminobenzidine (0.075 mg/ml; Sigma) and 0.04% nickel chloride as a chromogen. All washing steps were performed in TBS. BrdU- and Ki67-stained sections were mounted onto glass slides, cleared with Neo-Clear^®^ (Millipore) and cover-slipped using Neo-Mount^®^ (Millipore). BrdU and Ki67 positive cells were counted applying the simplified version of the optical fractionator principle as previously described (Kempermann et al, 1997) on every sixth section along the entire rostro-caudal axis of the dentate gyrus using a brightfield microscope (Leica DM 750). Synaptoporin-stained sections underwent a Nissl-staining before mounting them with Entellan (Merck). To prepare sections for Nissl staining, they were incubated for 20 min in each of the following solutions: staining buffer (4% sodium acetate, 0.96% acetic acid), followed by incubation in permeabilization solution (75% Ethanol, 0.025% Triton- X100) and staining buffer. Staining solution (0.1% cresyl violet in staining buffer) was applied for 20 min followed by differentiation of sections in 95% ethanol for 30 s and dehydration with isopropanol and xylene for 10 min each.

### Immunofluorescence staining

Immunofluorescent staining was performed for co-labeling of BrdU positive cells with NeuN and S100β as described (Steiner, Zurborg, Hörster, Fabel, & Kempermann, 2008). Briefly, sections were treated with 2 M HCl, washed extensively with PBS and blocked in PBS supplemented with 10% donkey serum and 0.2% Triton-X100 for 1 h at room temperature, followed by incubation with primary antibodies overnight at 4°C (rat anti-BrdU 1:500, Serotec; mouse anti-NeuN 1:100, Chemicon International; rabbit anti-S100β 1:200, Abcam). Secondary antibodies were incubated for 4 h at room temperature (anti-rat Alexa 488 1:500, anti-mouse Cy5 1:500, anti-rabbit Cy3 1:500; all from Jackson ImmunoResearch). Nuclei were counterstained using 4',6-diamidino-2-phenylindole (DAPI; 3.3 *μ*g/ml) for 10 min. All washing steps were performed in PBS. Sections were mounted onto glass slides and cover- slipped using Aqua-Poly/Mount (Polysciences, Inc.). Imaging was performed with the ZEISS Apotome and the Software AxioVision software with optical sectioning mode. To determine total numbers of new-born neurons and astrocytes, 100 randomly selected BrdU immuno- positive cells along the rostro-caudal axis of the dentate gyrus were investigated for coexpression with NeuN or S100β. The final numbers of surviving new neurons and astrocytes were obtained by multiplying the total number of BrdU positive cells (as determined by peroxidase-based immunohistochemistry) by the ratio of NeuN/BrdU-positive cells and S100β/BrdU-positive cells.

### Brain morphometry and volumetry

The mossy fiber (MF) projections are characterized by a high content of the presynaptic vesicle protein synaptoporin (Krebs et al., 2011; Singec et al., 2002), therefore the volumes of the MF projections were estimated on sections immunolabeled against synaptoporin and counterstained with Nissl for a better distinction between neuronal cell layers. Volumetric analysis was performed on every 6th section with a semiautomated morphometric system consisting of a CCD camera (Hitachi) connected to a light microscope (Leica DM-RXE) using a 10x objective and the Stereoinvestigator 7 software (MBF Bioscience). Structures were overlayed with the Cavalieri estimator probe grid of 25 *μ*m and every grid point belonging to the particular structure of interest was selected. Volume estimates were calculated in the software taking into account the sampling interval (240 *μ*m) and the section thickness (40 *μ*m).

For the analysis of the cortex thickness, the areas of motor, entorhinal and cingulate cortices were defined as described by (Diamond et al., 1964; 1985). We used the following coordinates of bregma: motor cortex −1.06 to −1.46, entorhinal cortex −2.30 to −2.80, cingulate cortex 1.34 to 0.50. Two to three constitutive sections from each animal were analyzed. Sections were scanned with a slide scanner (Axio Scan.Z1, Zeiss, Germany) and measured using the ZEN blue software (Zeiss, Germany).

### Analysis of blood samples

Blood was collected into EDTA-coated tubes (Sarstedt) from the abdominal cavity during the perfusion immediately after the right ventricle was opened. Blood samples were incubated for 1 to 2 h at room temperature, and centrifuged at 2000 × *g* for 15 min at 4 °C. Plasma was centrifuged a second time and stored at −80 °C. Plasma samples were assayed for glucose (Amplex red glucose/glucose oxidase assay kit, Invitrogen), cholesterol (Amplex red cholesterol assay kit, Invitrogen), triglycerides (Triglycerides colorimetric quantification kit, Abcam) and corticosterone (Corticosterone ELISA kit, Enzo) following the manufacturers’ instructions. Log-logistic concentration curves were calculated from standards in R using the drm function from the drc package (Ritz, Baty, Streibig, & Gerhard, 2015). Corticosterone and triglyceride measures were log-transformed to normality.

### Statisitics

All statistical analyses were carried out using the statistical software R (R Core Team, 2014). Data was tested for normality using the Shapiro-Wilk-test. For normally distributed measures, we used Welch’s Z-test to compare means and *F*-test to test for equality of variance between groups. For repeated measures (longitudinal data), a linear mixed regression was performed using the lmer function from the lme4 package (Bates, Mächler, Bolker, & Walker, 2015), and *p* values were obtained by the likelihood ratio test of the full model against the model without the analyzed effects. For non-normal data, we performed Wilcoxon-Rank-Sum test as a non-parametric analogue for the Z-test or Brown-Forsythe test as a more robust form of Levene’s test to compare the variances between groups. Differences were considered to be statistically significant at a *p* < 0.05. Graphs were generated using ggplot2 package (Wickham, 2011). Data is presented as means +- SEM.

All experiments were carried out with the experimenter blind to the experimental group.

## Acknowledgements

We thank Anne Karasinsky and Sandra Günther for taking care of our animals. We thank Alexander Garthe for his input while designing the behavioral experiments. We are grateful to all members of the Kempermann laboratory for assistance during the perfusion and collection of the various specimens.

---

**Supplementary Table 1.**
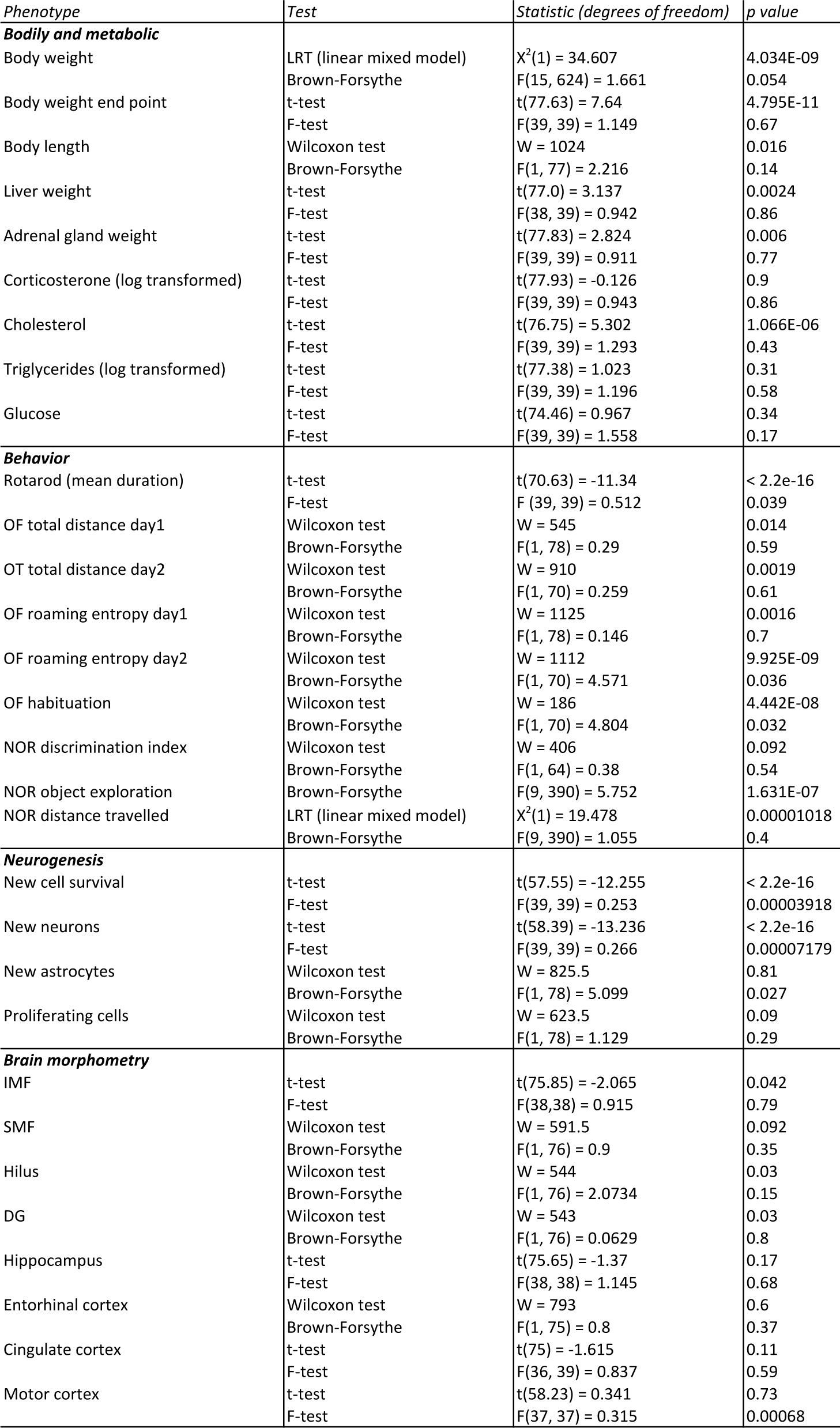
Test statistics and *p* values from tests comparing means and variances between CTRL and ENR mice. Abbreviations: LRT, likelihood ratio test; NOR, novel object recognition test; OF, open field test; IMF, infrapyramidal mossy fibers; SMF, suprapyramidal mossy fibers; DG, dentate gyrus.

## References

Bates, D., Mächler, M., Bolker, B., & Walker, S. (2015). Fitting Linear Mixed-Effects Models Using lme4. Journal of Statistical Software, 67(1), 1–48. http://doi.org/10.18637/jss.v067.i01

Bennett, E. L., Diamond, M. C., Krech, D., & Rosenzweig, M. R. (1964). Chemical and anatomical plasticity of brain. Science (New York, NY), 146(3644), 610–619.

Clemenson, G. D., Lee, S. W., Deng, W., Barrera, V. R., Iwamoto, K. S., Fanselow, M. S., & Gage, F. H. (2015). Enrichment rescues contextual discrimination deficit associated with immediate shock. Hippocampus, 25(3), 385–392. http://doi.org/10.1002/hipo.22380

Crusio, W. E., Schwegler, H., & van Abeelen, J. H. (1989). Behavioral responses to novelty and structural variation of the hippocampus in mice. I. Quantitative-genetic analysis of behavior in the open-field. Behavioural Brain Research, 32(1), 75–80.

Di Garbo, A., Mainardi, M., Chillemi, S., Maffei, L., & Caleo, M. (2011). Environmental enrichment modulates cortico-cortical interactions in the mouse. PLoS ONE, 6(9), e25285. http://doi.org/10.1371/journal.pone.0025285

Diamond, M. C. (2001). Response of the brain to enrichment. Anais Da Academia Brasileira De Ciencias, 73(2), 211–220.

Diamond, M. C., Johnson, R. E., Protti, A. M., & Ott, C. (1985). Plasticity in the 904-day-old male rat cerebral cortex. Experimental …, 87(2), 309–317. http://doi.org/10.1016/0014-4886(85)90221-3

Diamond, M. C., Krech, D., & Rosenzweig, M. R. (1964). The effects of an enriched environment on the histology of the rat cerebral cortex. The Journal of Comparative Neurology, 123, 111–120.

Diamond, M. C., Law, F., Rhodes, H., Lindner, B., Rosenzweig, M. R., Krech, D., & Bennett, E. L. (1966). Increases in cortical depth and glia numbers in rats subjected to enriched environment. The Journal of Comparative Neurology, 128(1), 117–126. http://doi.org/10.1002/cne.901280110

Fabricius, K., Helboe, L., Steiniger-Brach, B., Fink-Jensen, A., & Pakkenberg, B. (2010). Stereological brain volume changes in post-weaned socially isolated rats. Brain Research, 1345, 233–239. http://doi.org/10.1016/j.brainres.2010.05.040

Freund, J., Brandmaier, A. M., Lewejohann, L., Kirste, I., Kritzler, M., Kruger, A., et al. (2013). Emergence of Individuality in Genetically Identical Mice. Science (New York, NY), 340(6133), 756–759. http://doi.org/10.1126/science.1235294

Freund, J., Brandmaier, A. M., Lewejohann, L., Kirste, I., Kritzler, M., Kruger, A., et al. (2015). Association between exploratory activity and social individuality in genetically identical mice living in the same enriched environment. Neuroscience, 309(C), 140–152. http://doi.org/10.1016/j.neuroscience.2015.05.027

Garthe, A., Roeder, I., & Kempermann, G. (2015). Mice in an enriched environment learn more flexibly because of adult hippocampal neurogenesis. Hippocampus. http://doi.org/10.1002/hipo.22520

Hammelrath, L., Skokie, S., Khmelinskii, A., Hess, A., van der Knaap, N., Staring, M., et al. (2016). Morphological maturation of the mouse brain: An in vivo MRI and histology investigation. Neuroimage, 125, 144–152. http://doi.org/10.1016/j.neuroimage.2015.10.009

Jonker, R. M., Guenther, A., Engqvist, L., & Schmoll, T. (2013). Does systematic variation improve the reproducibility of animal experiments? Nature Methods, 10(5), 373–373. http://doi.org/10.1038/nmeth.2439

Kapogiannis, D., & Mattson, M. P. (2011). Disrupted energy metabolism and neuronal circuit dysfunction in cognitive impairment and Alzheimer’s disease. The Lancet Neurology, 10(2), 187–198. http://doi.org/10.1016/S1474-4422(10)70277-5

Kempermann, G., & Gage, F. H. (1999). Experience-dependent regulation of adult hippocampal neurogenesis: effects of long-term stimulation and stimulus withdrawal. Hippocampus, 9(3), 321–332. http://doi.org/10.1002/(SICI)1098-1063(1999)9:3<321::AID-HIP011>3.0.C0;2-C

Kempermann, G., Gast, D., & Gage, F. H. (2002). Neuroplasticity in old age: sustained fivefold induction of hippocampal neurogenesis by long-term environmental enrichment. Annals of Neurology, 52(2), 135–143. http://doi.org/10.1002/ana.10262

Kempermann, G., Kuhn, H. G., & Gage, F. H. (1997). More hippocampal neurons in adult mice living in an enriched environment. Nature, 386(6624), 493–495. http://doi.org/10.1038/386493a0

Krebs, J., Römer, B., Overall, R. W., Fabel, K., Babu, H., Brandt, M. D., et al. (2011). Adult Hippocampal Neurogenesis and Plasticity in the Infrapyramidal Bundle of the Mossy Fiber Projection: II. Genetic Covariation and Identification of Nos1 as Linking Candidate Gene. Frontiers in Neuroscience, 5, 106. http://doi.org/10.3389/fnins.2011.00106

Loseva, E., Yuan, T.-F., & Karnup, S. (2011). Genetics and human agency: comment on Dar-Nimrod and Heine (2011). Psychological Bulletin, 137(5), 825–828. http://doi.org/10.1037/a0024306

Melani, R., Chelini, G., Cenni, M. C., & Berardi, N. (2017). Enriched environment effects on remote object recognition memory. Neuroscience, 352, 296–305. http://doi.org/10.1016/j.neuroscience.2017.04.006

Miyauchi, M., Neugebauer, N. M., Oyamada, Y., & Meltzer, H. Y. (2016). Nicotinic receptors and lurasidone-mediated reversal of phencyclidine-induced deficit in novel object recognition. Behavioural Brain Research, 301, 204–212. http://doi.org/10.1016/j.bbr.2015.10.044

Mohammed, A. H., Zhu, S. W., Darmopil, S., Hjerling-Leffler, J., Ernfors, P., Winblad, B., et al. (2002). Environmental enrichment and the brain. Progress in Brain Research, 138, 109–133. http://doi.org/10.1016/S0079-6123(02)38074-9

Niell, C. M., & Stryker, M. P. (2010). Modulation of visual responses by behavioral state in mouse visual cortex. Neuron, 65(4), 472–479. http://doi.org/10.1016/j.neuron.2010.01.033

Nilsson, M., Perfilieva, E., Johansson, U., Orwar, O., & Eriksson, P. S. (1999). Enriched environment increases neurogenesis in the adult rat dentate gyrus and improves spatial memory. Journal of Neurobiology, 39(4), 569–578.

Nuzzo, R. (2014, February 13). Scientific method: statistical errors. Nature, pp. 150–152. http://doi.org/10.1038/506150a

Panza, F., Solfrizzi, V., Logroscino, G., Maggi, S., Santamato, A., Seripa, D., & Pilotto, A. (2012). Current epidemiological approaches to the metabolic-cognitive syndrome. Journal of Alzheimer’s Disease : JAD, 30 Suppl 2(s2), S31–75. http://doi.org/10.3233/JAD-2012-111496

R Core Team. (2014). R: A language and environment for statistical computing. Retrieved January 22, 2018, from http://www.R-project.org

Richter, S. H., Garner, J. P., Auer, C., Kunert, J., & Würbel, H. (2010). Systematic variation improves reproducibility of animal experiments. Nature Methods, 7(3), 167–168. http://doi.org/10.1038/nmeth0310-167

Richter, S. H., Garner, J. P., Zipser, B., Lewejohann, L., Sachser, N., Touma, C., et al. (2011). Effect of population heterogenization on the reproducibility of mouse behavior: a multilaboratory study. PLoS ONE, 6(1), e16461. http://doi.org/10.1371/journal.pone.0016461.

Ritz, C., Baty, F., Streibig, J. C., & Gerhard, D. (2015). Dose-Response Analysis Using R. PLoS ONE, 10(12), e0146021. http://doi.org/10.1371/journal.pone.0146021

Rosenzweig, M. R., & Bennett, E. L. (1996). Psychobiology of plasticity: effects of training and experience on brain and behavior. Behavioural Brain Research, 78(1), 57–65.

Romer, B., Krebs, J., Overall, R. W., Fabel, K., Babu, H., Overstreet Wadiche, L., et al. (2011). Adult hippocampal neurogenesis and plasticity in the infrapyramidal bundle of the mossy fiber projection: I. Co-regulation by activity. Frontiers in Neuroscience, 5, 107. http://doi.org/10.3389/fnins.2011.00107

Sale, A., Berardi, N., & Maffei, L. (2014). Environment and brain plasticity: towards an endogenous pharmacotherapy. Physiological Reviews, 94(1), 189–234. http://doi.org/10.1152/physrev.00036.2012

Schwegler, H., Lipp, H. P., Van der Loos, H., & Buselmaier, W. (1981). Individual hippocampal mossy fiber distribution in mice correlates with two-way avoidance performance. Science (New York, NY), 214(4522), 817–819.

Shemesh, Y., Sztainberg, Y., Forkosh, O., Shlapobersky, T., Chen, A., & Schneidman, E. (2013). High-order social interactions in groups of mice. eLife, 2, e00759. http://doi.org/10.7554/eLife.00759

Singec, I., Knoth, R., Ditter, M., Hagemeyer, C. E., Rosenbrock, H., Frotscher, M., & Volk, B. (2002). Synaptic vesicle protein synaptoporin is differently expressed by subpopulations of mouse hippocampal neurons. The Journal of Comparative Neurology, 452(2), 139–153. http://doi.org/10.1002/cne.10371

Steiner, B., Zurborg, S., Hörster, H., Fabel, K., & Kempermann, G. (2008). Differential 24 h responsiveness of Prox1-expressing precursor cells in adult hippocampal neurogenesis to physical activity, environmental enrichment, and kainic acid-induced seizures. Neuroscience, 154(2), 521–529. http://doi.org/10.1016/j.neuroscience.2008.04.023

Tashiro, A., Makino, H., & Gage, F. H. (2007). Experience-specific functional modification of the dentate gyrus through adult neurogenesis: a critical period during an immature stage. The Journal of Neuroscience : the Official Journal of the Society for Neuroscience, 27(12), 3252–3259. http://doi.org/10.1523/JNEUROSCI.4941-06.2007

Walsh, R. N., & Cummins, R. A. (1979). Changes in hippocampal neuronal nuclei in response to environmental stimulation. The International Journal of Neuroscience, 9(4), 209–212.

Wei, Y., Yang, C.-R., Wei, Y.-P., Ge, Z.-J., Zhao, Z.-A., Zhang, B., et al. (2015). Enriched environment-induced maternal weight loss reprograms metabolic gene expression in mouse offspring. The Journal of Biological Chemistry, 290(8), 4604–4619. http://doi.org/10.1074/jbc.M114.605642

Wickham, H. (2011). ggplot2. WIREs Comp Stat, 3, 180–185. http://doi.org/10.1002/wics.147

Winkler, A. M., Kochunov, P., Blangero, J., Almasy, L., Zilles, K., Fox, P. T., et al. (2010). Cortical thickness or grey matter volume? The importance of selecting the phenotype for imaging genetics studies. Neuroimage, 53(3), 1135–1146. http://doi.org/10.1016/j.neuroimage.2009.12.028

Wolfer, D. P., Litvin, O., Morf, S., Nitsch, R. M., Lipp, H.-P., & Würbel, H. (2004). Laboratory animal welfare: cage enrichment and mouse behaviour. Nature, 432(7019), 821–822. http://doi.org/10.1038/432821a

Wolfinger, R. D. (2013). Reanalysis of Richter et al. (2010) on reproducibility. Nature Methods, 10(5), 373–374. http://doi.org/10.1038/nmeth.2438

